# Multiple isotopic models using various kinetic fractionation coefficients to estimate *δ*^18^O of leaf water

**DOI:** 10.1101/403410

**Authors:** Yonge Zhang, Xinxiao Yu, Lihua Chen, Guodong Jia

## Abstract

Investigation of *δ*^18^O of leaf water may improve our understanding of the evapotranspiration partitioning and material exchange between the inside and outside of leaves. In this study, *δ*^18^O of bulk leaf water (*δ*_L,b_) was estimated by both isotopic–steady–state (ISS) and non–steady–state (NSS) assumptions considering the Péclet effect. Specifically, we carefully modified kinetic fractionation coefficients (*α*_k_). The results showed that the Péclet effect is required to predict *δ*_L,b_. On the diel time scale, both NSS assumption + Péclet effect (NSS + P) and ISS assumption + Péclet effect (ISS + P) using modified *α*_k_ (*α*_k–modified_) for *δ*_L,b_ showed a good agreement with observed *δ*_L,b_ (*p* > 0.05). When using previously proposed *α*_k_, however, both NSS + P and ISS + P were not reliable estimators of *δ*_L,b_ (*p* < 0.05). On a longer time scale (days), estimates of daily mean *δ*_L,b_ from ISS + P outperformed the estimates from NSS + P when using the same *α*_k_ values. Also, the employment of *α*_k–modified_ improved model performance in predicting daily mean *δ*_L,b_ compared to the use of previously proposed *α*_k_. Clearly, special care must be taken concerning *α*_k_ when using isotopic models to estimate *δ*_L,b_.

**Highlight:** For hourly and daily mean data sets, the employment of modified kinetic fractionation coefficients significantly improved model performance for *δ*^18^O of bulk leaf water.

## Instruction

Oxygen isotope compositions of leaf water indicate material exchange between the inside and outside of leaves. For example, *δ*^18^O of atmospheric CO_2_ and atmospheric O_2_ are partly dependent on the *δ*^18^O of leaf water (Hoffmann *et al*., 2004; Luz and Barkan, 2011; Welp *et al*., 2011). The oxygen isotope signal of leaf water can be incorporated in carbohydrates, which can be retained in cellulose (Sullivan *et al*., 2006; Barbour, 2007; Gessler *et al*., 2014). Therefore, the oxygen isotope signature of leaf water plays an important role in constraining models of the global carbon cycle and reconstructing past climates. In addition, the investigation of leaf water isotope composition has significant implications for hydrology. In recent years, leaf isotopic models have been developed and improved for the partitioning of evapotranspiration (Sun *et al*., 2014; Wang *et al*., 2015). However, these multiple uses require a thorough description and prediction of leaf water isotope compositions.

Isotope fractionation does not occur during water absorption and transport before transpiration, except in saline plants (Walker *et al*., 2001), resulting in *δ*^18^O values of water in non-transpiring plant organs (i.e. twigs) similar to those of the source water (*δ*_source_). However, transpiration results in isotopic enrichment at evaporating sites within leaves (Cernusak *et al*., 2016). According to Sun *et al*. (2014), H_2_^18^O is enriched at the evaporation sites of the stomata during water vapor diffusion from leaves to the ambient air as a result of isotope fractionation effects, namely temperature–controlled equilibrium isotope effects (equilibrium fractionation coefficient, *α*^+^) and diffusion–controlled kinetic isotope effects (kinetic fractionation coefficient, *α*_k_) (Luo *et al*., 2013). The isotopically enriched water can then diffuse away from the evaporative sites into other parts of the leaf. Therefore, the *δ*^18^O of leaf water becomes unevenly distributed, and the *δ*^18^O of bulk leaf water (*δ*_L,b_) is depleted relative to the *δ*^18^O of leaf water at the evaporative sites (*δ*_L,e_) (Farquhar and Cernusak, 2005; Farquhar *et al*., 2007; Song *et al*., 2013).

Gonfiantini *et al*. (1965) firstly discovered leaf water enrichment because of the evaporative process of transpiration. In the same year, Craig & Gordon (1965) proposed a model to describe the isotopic enrichment of open water bodies during evaporation, and this model can be modified to be applied in leaf water enrichment (Dongmann *et al*., 1974). Recently, the traditional isotopic–steady–state (ISS) assumption (i.e., modified Craig–Gordon model), used to estimate *δ*_L,e_, has been challenged widely (Farquhar and Cernusak, 2005; Sun *et al*. 2014). The ISS assumption requires that *δ*^18^O of transpiration (*δ*_T_) is equal to *δ*_source_ (Yepez *et al*., 2003; Wang *et al*., 2015). Indeed, this assumption can only be approximately satisfied when transpiration is intensive (usually during midday) or over a long time scale (Wen *et al*. 2008). Thus, a more complex non–steady–state (NSS) assumption, considering heavy water fractionation during transpiration, has been introduced to accurately predict diel variations in *δ*_L,e_ (Farquhar and Cernusak, 2005). Moreover, studies have consistently suggested that the discrepancy between leaf water enrichment at the evaporative sites beyond the source water (Δ_l,,e_) and bulk leaf water enrichment beyond the source water (Δ_l,,b_) is due to the Péclet effect (*P*) (Farquhar *et al*., 2003; Farquhar and Cernusak, 2005; Song *et al*. 2013). This effect means that unenriched source water in leaf veins, which is transferred to evaporative sites, is opposed by a backwards diffusion of enriched water from the evaporative sites to the leaf veins, which holds in both ISS and NSS and can be combined to predict leaf water enrichment (Farquhar *et al*., 2003; Farquhar and Cernusak, 2005). Various models have been introduced; however, it remains unclear which one is the best to describe leaf water enrichment over time under field conditions.

Furthermore, H_2_ enrichment is predominately influenced by equilibrium isotope effects, whereas ^18^O enrichment is predominately influenced by the kinetic fractionation coefficient (Cernusak *et al*., 2016). Equilibrium isotope effects are dependent on temperature and have been well described and understood, while kinetic isotope effects are more complex. Kinetic fractionation occurs because the heavier isotopes diffuse more slowly than the lighter ones, which include resistance associated with water vapor transport and molecular diffusion (Dubbert *et al*., 2013; Zhang *et al*., 2018). The adequate determination of *α*_k_ is still subject of debate, and both values previously assigned are still in use (Rothfuss *et al*., 2012; Dubbert *et al*., 2013). According to Merlivat (1978), the molecular diffusion coefficient D*v*/D^i^*v*, ratio of the diffusivity of H_2_^18^O and H_2_O in air equals 1.0285. However, recently, Cappa *et al*. (2003) have claimed that this value needs to be revised and determined a new value of 1.032 on the basis of kinetic theory. Therefore, most studies have used an approximate value of 1.032 (*α*_k–Cappa_) or 1.0285 (*α*_k–Merlivat_), although under field conditions, *α*_k_ is variable (Rothfuss *et al*., 2012). In contrast to expressing *α*_k_ by pure molecular diffusion, researchers associate *α*_k_ with the fractionation of pure molecular diffusion by an exponent *nk*, which is expressed as *α*_k_ = (*Dv*/*D*^i^*v*)^*nk*^ (Mathieu and Bariac, 1996; Stewart, 1975; Dubbert *et al*., 2013). The exponent *nk* is dominated by the nature of the transport (Stewart 1975). For soil water evaporation, Dubbert *et al*. (2013) have linked *nk* with the soil water content, thereby reasonably well predicting the ^18^O enrichment of soil evaporation water. However, studies on including various resistance influences on the kinetic effects for predicting leaf water enrichment are still scarce.

The estimation of *δ*^18^O of leaf water largely depends on the leaf isotopic models and kinetic effects. In this study, *δ*_L,b_ values were simulated by ISS assumption with the Péclet effect (ISS+ P) and NSS assumption with the Péclet effect (NSS + P). Specifically, we carefully modified *α*_k_ (*α*_k–modified_) and tested different model performances when using *α*_k–modified_ and previously proposed *α*_k_ values (*α*_k–Cappa_ and *α*_k–Merlivat_). Our objectives were (1) to determine which model is useful to predict *δ*^18^O of bulk leaf water and (2) to examine the validity of *α*_k–modified_ for the estimation of *δ*^18^O of bulk leaf water.

### Theoretical background

ISS assumption and kinetic fractionation coefficient

The *δ*^18^O of leaf water at evaporative sites under ISS conditions, *δ*_L,e–ISS_, is commonly expressed in terms of kinetic fractionation factors *ε*_k_ (‰), equilibrium fractionation factors *ε*^+^ (‰), *δ*^18^O of xylem water *δ*_X_ (‰), *δ*^18^O of water vapor outside the leaf *δ*_v_ (‰), and the relative humidity of air outside the leaf relative to leaf temperature *h*_v_ (%). This way, *δ*_L,e–ISS_ can be calculated as follows (Farquhar and Cernusak, 2005; Kahmen *et al*., 2008):

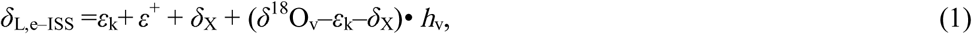

where 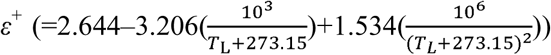 changes as a function of leaf temperature *T*_L_ (°C) (Sun *et al*., 2013), *h*_v_ is the ratio of water vapor pressure outside the leaf (0.611×exp(17.27×*T*_a_/(*T*_a_+ 273.1))×*h*_a_, with *T*_a_ and *h*_a_ being air temperature in °C and relative humidity in %, respectively) to saturated water vapor pressure in the leaf intercellular space (0.611×exp(17.27×T_L_/(T_L_+273.15))); *ε*_k_ is given by (Luo *et al*.,(2)13):

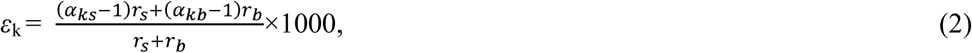

where *r*_s_ is stomatal resistance (m^2^•s•mol^−1^), *r*_b_ is the boundary layer resistance (m^2^•s•mol^−1^), *α*_ks_ is the kinetic fractionation coefficient associated with water vapor diffusion through the stoma (m^2^•s•mol^−1^), *α*_kb_ is the kinetic fractionation coefficient associated with water vapor diffusion through the boundary layer. As *r*_b_ is far smaller than *r*_s_, Equation (2) is rearranged as follows:

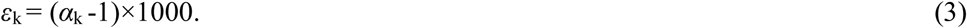

Considering pure molecular diffusion, the *α*_k_ value is generally 1.032 (Cappa *et al*., 2003) or 1.0285 (Merlivat, 1978); therefore, *ε*_k_is equivalent to 32‰ (*ε*_k–Cappa_) or 28‰ (*ε*_k–Merlivat_). Deviating from the theory, Stewart (1975) presented *α*_k_ as follows:

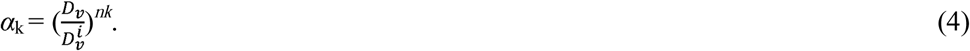

As field conditions generally vary between diffusive and laminar, for simulation isotope profiles of soil evaporating water, Mathieu and Bariac (1996) empirically expressed the contribution rate of turbulent resistance to total transport resistance in terms of soil water content. In addition, a recent study has suggested that the influence of aerodynamic resistance on kinetic effects needs to be considered (Lee *et al*., 2009). This study did not take into account aerodynamic resistance as it is insignificant or extremely low in dense forests with relatively low wind speeds and high canopy height. Similar to the calculation of kinetic fractionation coefficients, which empirically linked the exponent *nk* with soil water content, we related the exponent *nk* to leaf surface water content *θ*_surf_ (%) as well as saturated *θ*_sat_ (%) and residual leaf water content *θ*_r_ (%) to describe the nature of *α*_k_ associated with transpiration. In this sense, the exponent *nk*, associated with transpiration, is defined as follows:

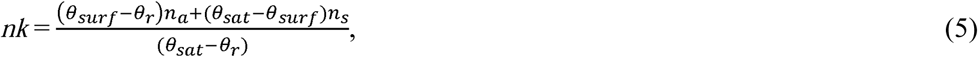

where *n*_a_ (=0.5) and *n*_s_ (=1) are constants; θ_surf_, θ_sat_, and θ_r_ are expressed as follows:

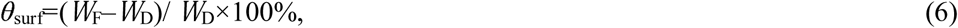

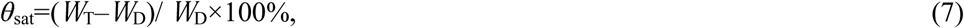

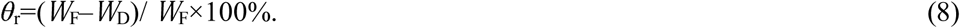

where *W*_F_, W_T_, and *W*_D_ are leaf fresh weight (*g*), turgid weight (*g*), and dry weight(*g*), respectively. In this study, θ_surf_ varied with little diel amplitudes (data not shown), so we considered θ_surf_ as constant on diel time scales to avoid an unnecessary complexity that does not significantly improve the model performance. Moreover, the differences in θ_surf_ among different days during the dry season or the wet season were not significant, whereas θ_surf_ differed between dry and wet season, with values of 107.7% for the dry season and 116.1% for the wet season. The values of θ_r_ were 51.8 and 53.7% for the dry season and the wet season, respectively, while θ_sat_ values for both seasons were 145%. Therefore, the calculated *nk* was 0.66 for the dry season and 0.70 for the wet season, and *α*_k_ was modified (*α*_k–modified_) as 1.0187 for the dry season and 1.0199 for the wet season.

The Péclet effect (*P*)

For leaves, researchers have described the Péclet effect in terms of leaf transpiration rate *T*_r_ (mol•m^−2^•s^−1^), the molar density of water *C* (= 55.56 × 10^3^ mol•m^−3^), the diffusion coefficient of H_2_^18^O in water *D*, and the effective path length for the effect *L* (m) (Sullivan *et al*., 2007). Therefore, the Péclet effect (*P*) can be described as follows:

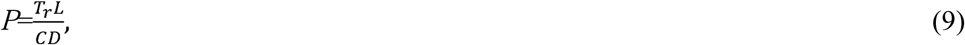

where *C* (=55.56×10^3^ mol•m^−3^) is constant and *D* (= 119 × 10^−9^ × exp (–637/(136.15 + T_L_))) is temperature–dependent. The direct measurement of *L* is still difficult, but recent studies have shown that variations of *L* could be explained by variations in *T*_r_ (Zhou *et al*., 2011; Ferrio *et al*., 2012). Our study calculated *L* on the basis of *L*–*T*_r_ dynamics as proposed by Song *et al*. (2013), who carried out research on six field-grown broad–leaf and coniferous tree species and found that *L* values are significantly and negatively correlated with 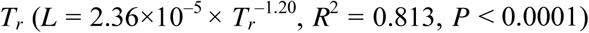.

Under ISS conditions, bulk leaf water enrichment beyond the source water, Δ_L,b–ISS_ (= (*δ*_L,b–ISS_– *δ*_X_)/(1+*δ*_X_), ‰), can be modeled by leaf water enrichment at the evaporative sites beyond the source water, Δ_L,e–ISS_(=(*δ*_L,e–ISS_–*δ*_X_)/(1+*δ*_X_), ‰) (Farquhar and Lloyd., 1993). In such a condition, Δ_L,b–ISS_ is expressed as follows:

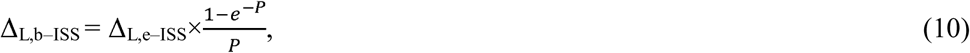

i.e.,

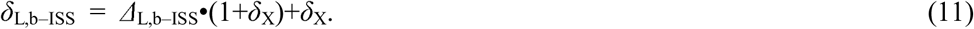

The NSS assumption

When stomatal conductance is low and/or when leaf water concentrations are high, NSS effects on the isotopic enrichment of leaf water should not be neglected (Cernusak *et al*., 2016). Under NSS conditions, leaf water enrichment at the evaporative sites beyond the source water, Δ_L,e–NSS_ (‰), can be described by the improved model developed by Farquhar and Cernusak (2005) on the basis of Dongmann *et al*. (1974):

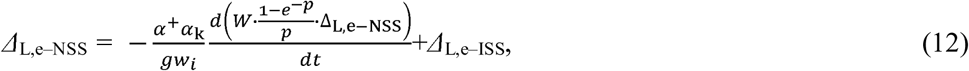

where *α*^+^is 1 + *ε*^+^, *g* is total leaf conductance (mol•m^−2^•s^−1^), *w*_i_ (= *W*_a_/*h*_v_, with *W*_a_ referring to the mole fraction of water vapor outside the leaf in mol•mol^−1^) is the mole fraction of water vapor in the leaf intercellular spaces (mol•mol^−1^), *W* is leaf water concentration (mol•m^−2^), and *t* is time (s). Note that Δ_L,e–NSS_ appears on both the right and the left side, making it difficult to calculate Δ_L,e–NSS_. To avoid iteratively calculating Δ_L,e–NSS_, Kahmen *et al*. (2008) advised a simpler approach to calculate Δ_L,e–NSS_. Over a relatively short time span,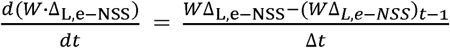,and Equation (12) changes as follows (Kahmen *et al*., 2008):

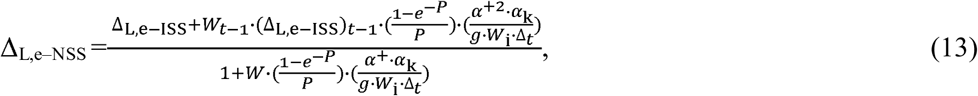

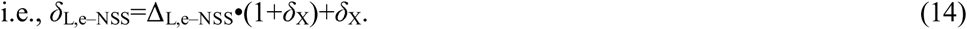

Assuming NSS conditions, bulk leaf water enrichment beyond the source water, Δ_L,b–NSS_, is expressed as follows (Farquhar and Cernusak, 2005):

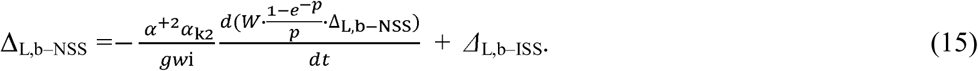

The calculation of Δ_L,b–NSS_ is similar to that of Δ_L,e–NSS_. Thus, Equation (15) is transformed in the following formula (Kahmen *et al*., 2008):

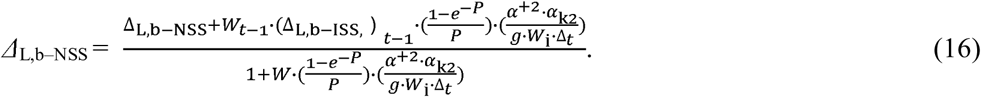

In addition,

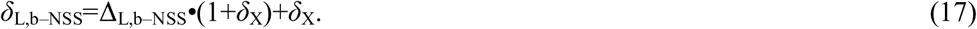

## Material and methods

### Study site

The experiment was conducted in 2017 at the Jiufeng National Forest Experimental Station (116°05É, N40°03´N). The station, at an elevation of 100 ∼ 1,153 m above sea level, is situated in Beijing in a rocky mountainous area of north China. The climate is a typical continental monsoon climate characterized by four distinct seasons. The growing season in this area typically stretches from April to October. Mean annual precipitation is about 650 mm, with an uneven temporal distribution; 70 ∼ 80% occur between July and September. Therefore, the period from April to June is the dry season, while the period from July to September is the rainy season. Mean annual potential evaporation is between 1,800 and 2,000 mm, which is significantly higher than precipitation. Mean annual temperature is 11.6°C, and mean monthly temperature ranges between –5 and 30°C, with maximum values in July and minimum values in November.

At the experimental station, the main tree species are drought–tolerant coniferous species. The central area (about 1 km^2^) of the sampling site, at an elevation of about 250 m, is dominated by 50 ∼ 60 years old *Platycladus orientalis* trees with a similar growth status; average tree height is 8 m and average diameter at breast height 20 cm. The forest crown density is about 0.8.

### Gas exchange and leaf and xylem water isotopic measurement

The experiment was carried out in 2017 at three typical sunny days in the dry season (May 24, May 30 and June 5,) and three typical sunny days in the wet season (July 29, August 12 and August 17). Diel variations in leaf transpiration rate (*T*_r_), stomatal conductance (*g*_s_), and leaf temperature (*T*_L_) were determined every hour from 0:00 to 24:00 using a portable photosynthesis system (LI–COR. USA, LI–6400). Gas exchange measurements were conducted on at least the top three intact mature leaves of the canopy from three *Platycladus orientalis* trees. Subsequently, these conifer leaves were sampled, covered with foil, and kept in liquid nitrogen tanks. In addition, annual branches from the upper canopy were collected from each of the three standard sample trees (three samples/tree), stored in 50–ml centrifuge tubes, tightly sealed, and stored in a portable refrigerator.

To determine bulk leaf water and xylem water, primary veins of the leaves were removed and the bark was removed from the branches. Subsequently, a cryogenic vacuum distillation system (Los Gatos Research Inc. USA, Li–2100) was used to extract leaf and xylem water. After extraction, *δ*^18^O values of bulk leaf water (*δ*_L,b–observed_) and xylem water were analyzed using a liquid water isotope analyzer (Los Gatos Research Inc. USA, DLT–100) with a measurement precision of ± 0.1‰.

### Leaf water concentration and leaf water content measurement

On each sampling day, intact leaves from the upper canopy of each sampling tree were detached from twigs hourly from 0:00 to 24:00 and immediately weighed on an electronic scale to determine leaf fresh weight (*FW*_L_). Image–processing software for Photoshop was used for leaf area (*LA*) measurement (Xiao *et al*. 2005). Thereafter, these leaves were soaked in deionized water for 24 h and then weighted again to determine leaf turgid weight (*TW*_L_). Finally, all leaves were dried at 80°C for 48 h and weighted to obtain dry weight (*DW*_L_); leaf water concentration (*W*) was calculated on the basis of leaf weight and leaf area as *W* = (*FW*_L_–*DW*_L_)/*LA*) (Kahmen *et al*. 2008).

### Micrometeorological and atmospheric water vapor measurement

The sampling site was equipped with an 18–m high meteorological tower to continuously output micrometeorological data at 10–minute intervals. A temperature–humidity compound sensor (Onset. USA, HOBO–U30), installed at a height of 8 m, was used to measure air temperature (*T*_a_) and relative humidity (*h*_a_) at the upper canopy layer. In addition, a water vapor isotope analyzer (Los Gatos Research Inc. USA, DLT–100), which is based on the Off–Axis Integrated Cavity Output Spectroscopy (OA–ICOS) approach, with a measurement precision of ± 0.2‰ and a data output frequency of 1 Hz, was used to measure atmospheric water vapor concentration (*W*_a_) and *δ*^18^O (*δ*_v_) at the upper canopy layer. Each measurement at the upper canopy lasted for 10 minutes. After measuring atmospheric water vapor three times, an extra 30 minutes were required to measure the standard water vapor concentration and *δ*^18^O for calibration of the measured *δ*_v_. In total, a full measurement cycle lasted for 1 hour.

### Data analysis

Differences between observed and modeled *δ*_L,b_ as well as between observed *δ*_L,b_ and modeled *δ*_L,e_ were analyzed by one–way analysis of variance (ANOVA). Differences in modeled *δ*_L,b_ and *δ*_L,e_, calculated by assuming different model assumptions and using various *α*_k_ values, were also analyzed by one–way ANOVA. Results were considered statistically significant at *p* < 0.05. Deviations caused by modeled *δ*_L,b_ and *δ*_L,e_ from observed *δ*_L,b_ were absolute differences between observed and modeled *δ*_L,b_ as well as between observed *δ*_L,b_ and modeled *δ*_L,e_, respectively. The presented values are means ± standard deviations (SD).

## Results

### 3.1 Environmental factors and atmospheric water vapor *δ*^18^O

On diel time scales, *T*_a_ represented an inverted-V shape, reaching maximum values during 12:00 ∼ 16:00 and decreasing to minimum values at night, with diel amplitudes between 7.09 and 15.02 □ (Fig. 1). The *h*_a_ showed an opposite trend to that of *T*_a_ on diel time scales, with diel amplitudes between 23.56 and 39.44%. The average *T*_a_ and *h*_a_ values in the dry season were 23.30□ and 45.50%, respective; while the average values in the wet season were 28.30□ and 65.03%, respectively (Fig. 1).

**Figure 1.**
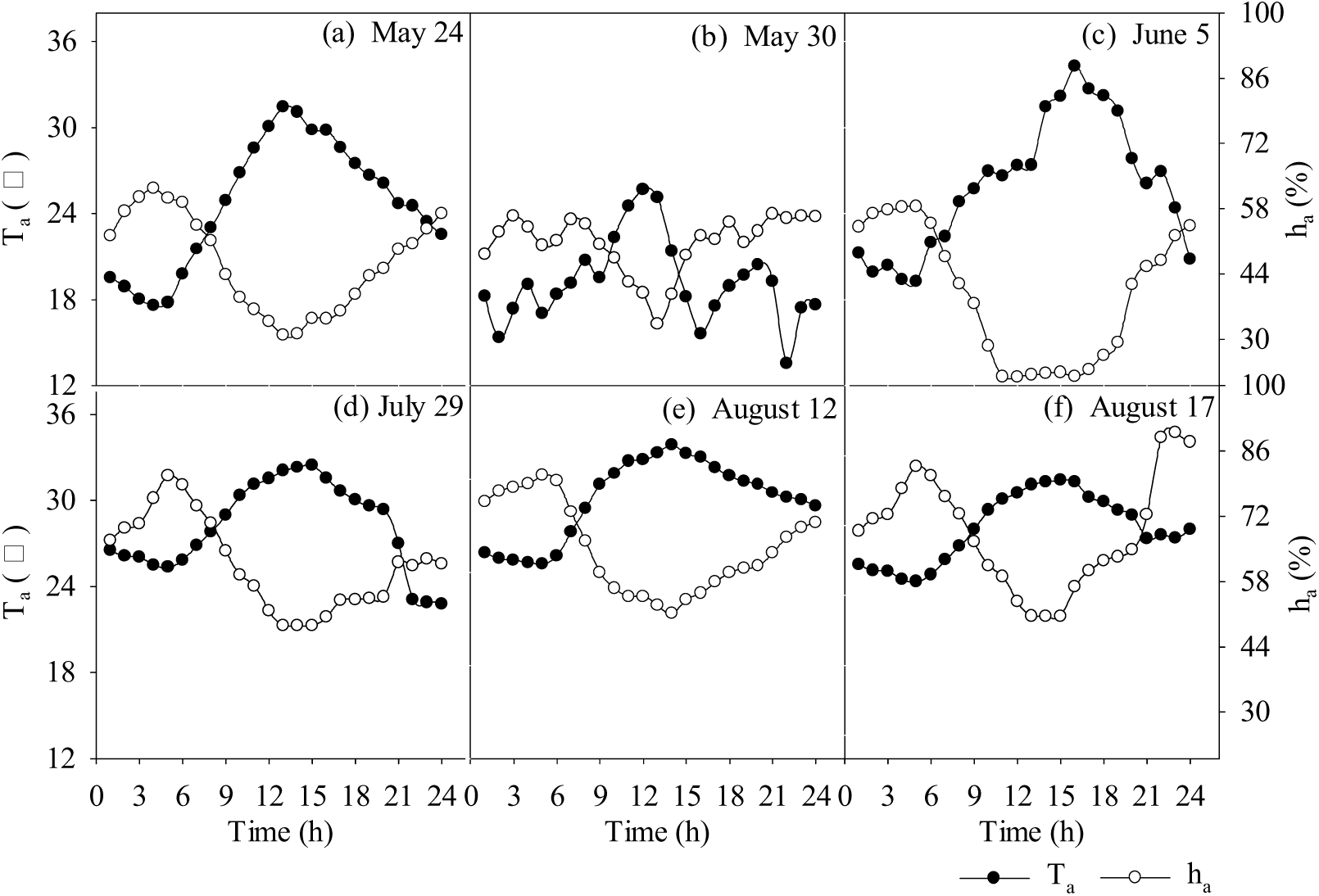
Variations in air temperature (*T*_a_) and relative humidity (*h*_a_) during the six experimental days.

However, *W*_a_ and *δ*_v_ varied with no clear diel patterns (Fig. 2). The *W*_a_ in the wet season (20.50 to 32.58 mmol•mol^−1^) was higher than that in the dry season (9.67 to 19.73 mmol•mol^−1^), while *δ*_v_ in the wet season (–19.27 to –13.11‰) was generally lower than that in the dry season (–17.73 to –10.00‰) (Fig. 2).

**Figure 2.**
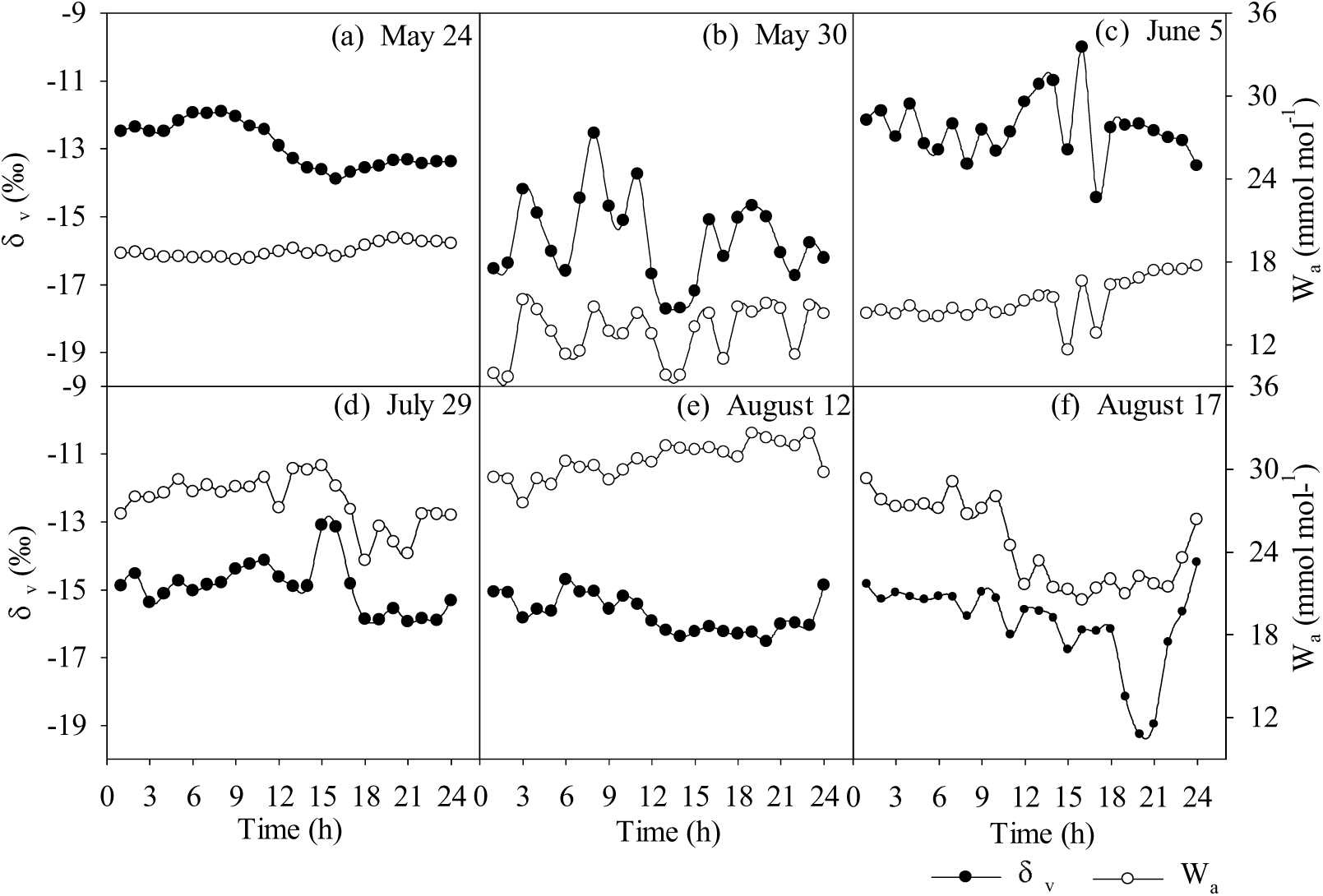
Variations in atmospheric water vapor concentration (*W*_a_) and *δ* O (*δ*_v_) during the six experimental days.

### 3.2 Diel variations in *T*_r_ and *g*_s_

Diel variations in leaf water enrichment were highly correlated with *T*_r_ and *g*_s_. On diel time scales, *T*_r_ and *g*_s_ showed similar patterns of variability (Fig. 3). The greatest *T*_r_ and *g*_s_ values were recorded around midday (11:00 ∼ 13:00), and their lowest values, approaching zero, were reached during the dark period. The *T*_r_ and g_s_ values in the dry season ranged from 0 to 0.15 mmol•m^−2^•s^−1^ and from 0.01 to 1.67 mol•m^−2^•s^−1^ and were, in general, lower than the values in the wet season, which ranged from 0.01 to 0.20 and from 0.01 to 2.25 mol•m^−2^•s^−1^, respectively. Midday depression of *T*_r_ and *g*_s_ was observed around 14:00 in the dry season, while no such phenomenon occurred in the wet season (Fig. 3).

**Figure 3.**
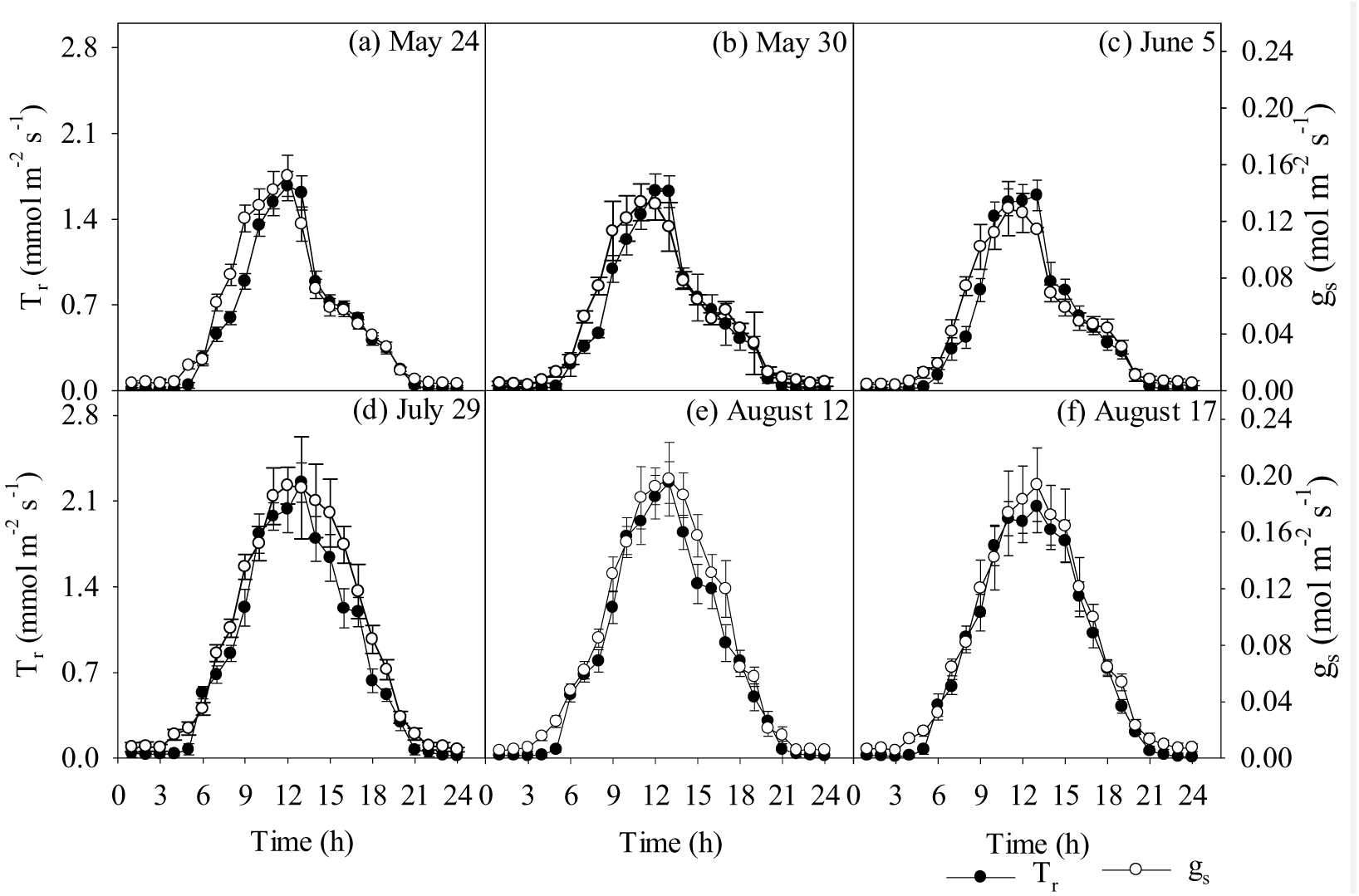
Diel variability of leaf transpiration rate (*T*_r_) and stomatal conductance (*g*_s_) over the sampling days. Data represent mean values ± SD.

### 3.3 *δ*^18^O of xylem water

The *δ*_X_ varied with no clear diel trend, and the diel amplitude value was between 0.43 and 2.73‰ (Fig. 4), indicating that the trees used identical or similar source water during the day. The *δ*_X_ values in the dry season varied from –7.80 to –4.38‰ and were thus lower than those in the wet season, which varied from –3.76 to –1.04 ‰ (Fig. 4).

**Figure 4.**
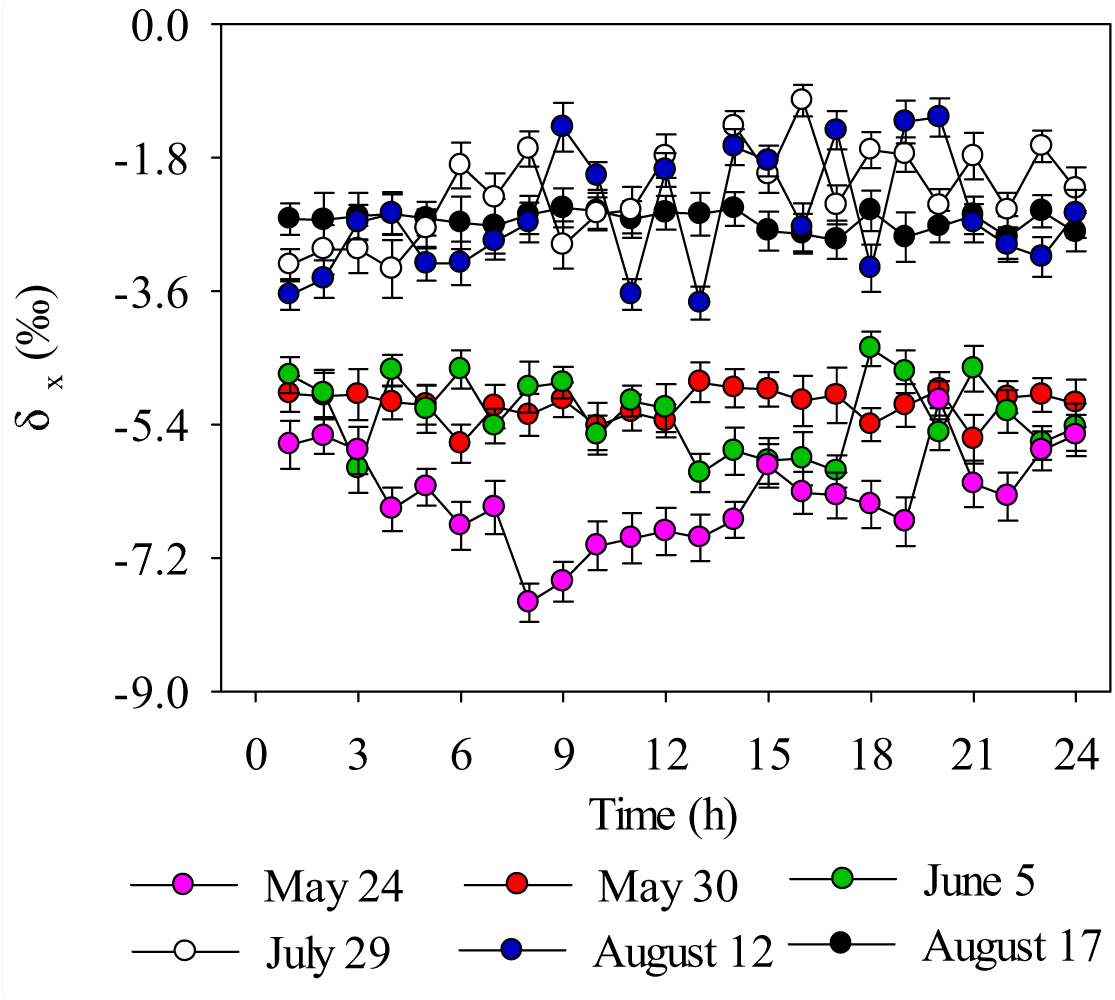
Diel variation of *δ*^18^O of xylem water (*δ*_X_). The *δ*_X_ are represented as mean values ± SD.

### 3.4 *δ*^18^O of bulk leaf water

The diel amplitude of observed *δ*_L,b_ (*δ*_L,b–observed_) ranged between 6.91 and 10.73‰ (Fig. 5), reaching maximum values during midday and minimum values in the early morning or evening. In the dry season, *δ*_L,b–observed_ ranged from 2.96 to 13.21‰. In contrast, *δ*_L,b–observed_ in the wet season ranged between −3.11 and 9.92‰.

**Figure 5.**
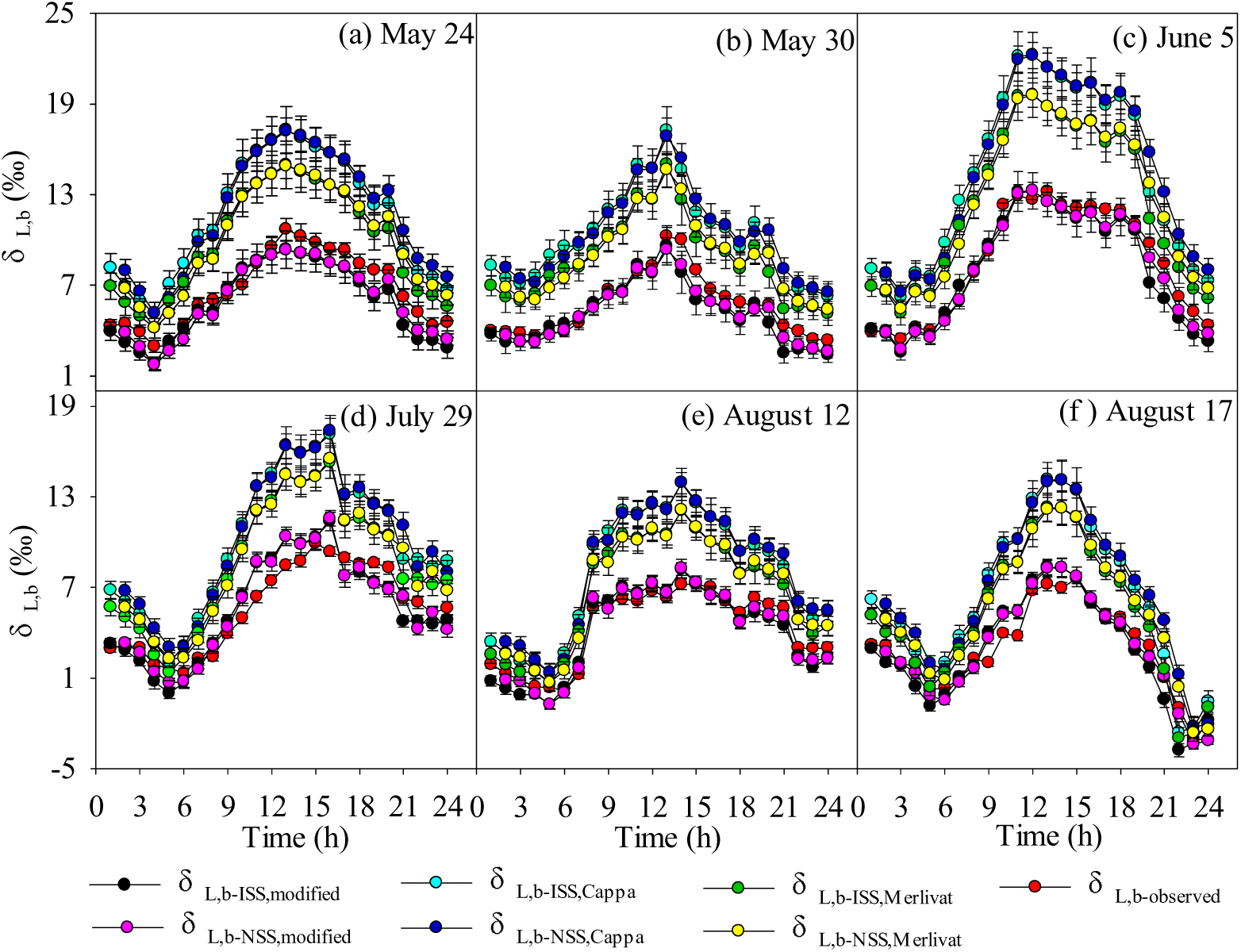
Diel variations in observed and modeled *δ*^18^O of bulk leaf water (*δ*_L,b_). *δ*_L,b–observed_ is observed *δ*_L,b_; *δ*_L,b–ISS,Cappa_ and *δ*_L,b–ISS,Merlivat_ are simulated by ISS assumption (Eqn. 11) using *α*_k2–Cappa_ and *α*_k2–Merlivat_, respectively; *δ*_L,b–ISS,modified_ is simulated by ISS assumption (Eqn. 11) using *α*_k2–modified_; *δ*_L,b–NSS,Cappa_ and *δ*_L,b–NSS,Merlivat_ are simulated by NSS assumption (Eqn. 17) using *α*_k2–Cappa_ and *α*_k2–Merlivat_, respectively; *δ*_L,b–NSS,modified_ is simulated by NSS assumption (Eqn. 17) using *α*_k2–modified_. The modeled and observed *δ*_L,b_ are represented as mean values ± SD. Data are mean values ± SD.

Diel variations in modeled *δ*_L,b_ are also shown in Figure 5. Considering the Péclet effects, the *δ*_L,b_ were modeled assuming both ISS and NSS conditions and using different formulations for *α*_k_ values. Using *α*_k–modified_, both the ISS (*δ*_L,b–NSS,modified_, –3.39 ∼ 13.25‰) and NSS assumptions for *δ*_L,b_ (*δ*_L,b–ISS,modified_, –3.77 ∼ 13.37‰) showed a good agreement with *δ*_L,b–observed_ (–3.11 ∼ 13.21‰). While using *α*_k–Merlivat_, modeled values of *δ*_L,b_ assuming the ISS (*δ*_L,b–ISS,Merlivat_, –2.97 ∼ 19.60‰) and NSS conditions (*δ*_L,b–NSS,Merlivat_, –2.61 ∼ 19.58‰) tended to overestimate *δ*_L,b_. In contrast, prediction of *δ*_L,b_ using *α*_k2–Cappa_, assuming both ISS (*δ*_L,b–ISS,Cappa_, –2.60 ∼ 22.23‰) and NSS conditions (*δ*_L,b–NSS,Cappa_, –2.24 ∼ 22.21‰), led to worst results than those one would calculate using *α*_k–modified_ or *α*_k–Merlivat_ (Fig. 5).

### 3.5 *δ*^18^O of leaf water at the evaporative sites

The diel variations in modeled *δ*_L,e_, assuming both ISS and NSS conditions and using different formulations to determine *α*_k_ values, were in accordance with the approximately dome–shaped diel patterns of modeled *δ*_L,b_ (Fig. 6). Values of *δ*_L,e–ISS,Cappa_ and *δ*_L,e–NSS,Cappa_ ranged from –2.99 to 24.27‰ and from –2.62 to 23.94‰, respectively, while *δ*_L,e–ISS,Merlivat_ and *δ*_L,e–NSS,Merlivat_ ranged from –2.55 to 27.39‰ and from –2.19 to 27.04‰, respectively. In contrast, *δ*_L,e–ISS,modified_ and *δ*_L,e–NSS,modified_ were characterized at –3.94 ∼ 16.74‰ and –3.55 ∼ 16.50‰. Clearly, the modeled *δ*_L,e_ values by using *α*_k2–Merlivat_ and *α*_k2–Cappa_ were significantly greater than those one would calculate using *α*_k–modified_ (*p <* 0.05; Fig. 6).

**Figure 6.**
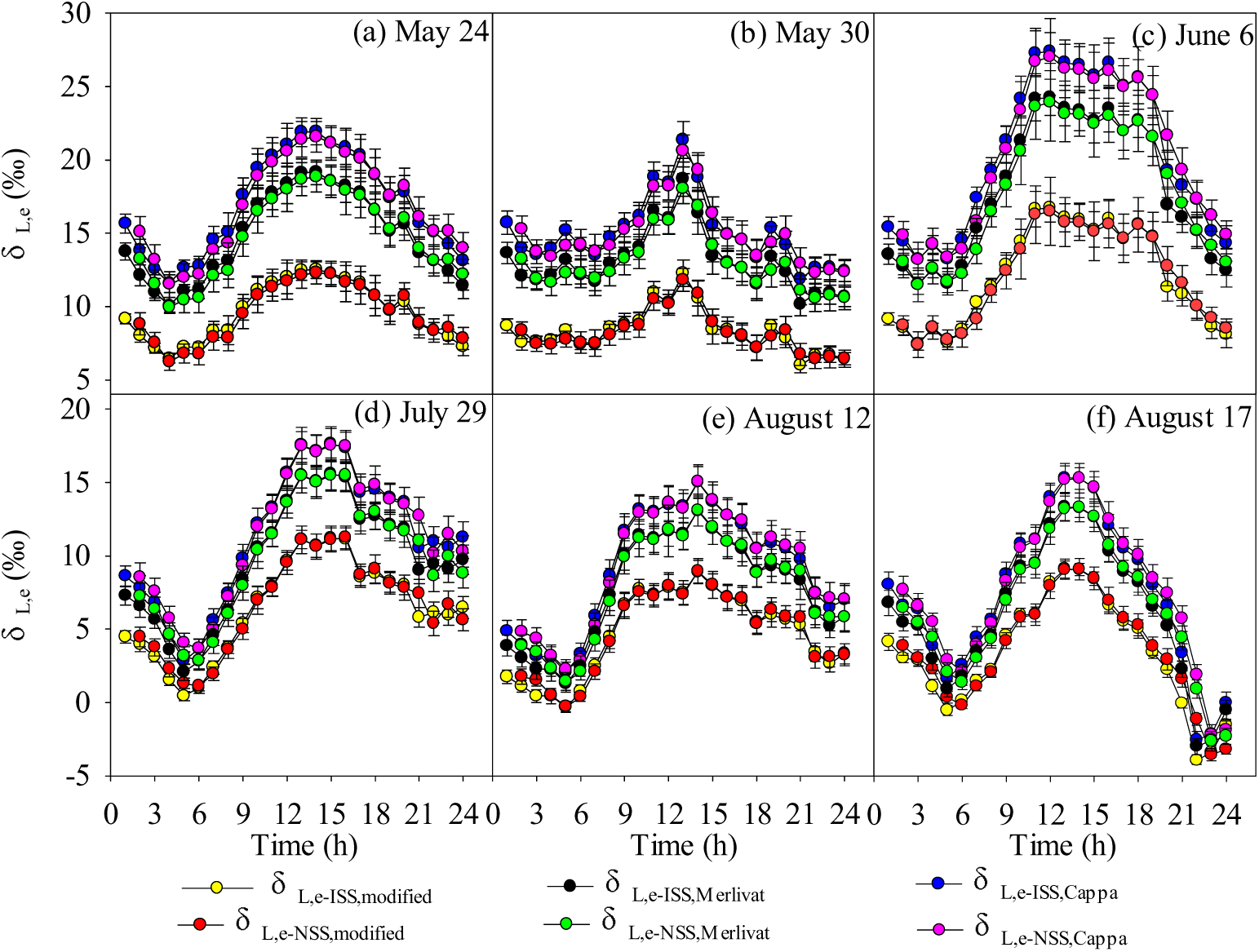
Diel variation in modeled *δ*^18^O of leaf water at evaporative sites (*δ*_L,e_). *δ*_L,e–ISS,Cappa_ and *δ*_L,e–ISS,Merlivat_ are simulated by ISS assumption (Eqn. 1) using *α*_k2–Cappa_ and *α*_k2–Merlivat_, respectively; *δ*_L,e–ISS,modified_ is simulated by ISS assumption (Eqn. 1) using *α*_k2–modified_; *δ*_L,e–NSS,Cappa_ and *δ*_L,e–NSS,Merlivat_ are simulated by NSS assumption (Eqn. 14) using *α*_k2–Cappa_ and *α*_k2–Merlivat_, respectively; *δ*_L,e–NSS,modified_ is simulated by ISS assumption (Eqn. 14) using *α*_k2–modified_. The modeled *δ*_L,e_ are represented as mean values ± SD.

## Discussion

*δ***_L,b_ estimated by different assumptions and** *α***_k_ values**

In this study, *δ*_X_ in the wet season was significantly greater than that in the dry season (Fig. 4), indicating that *P. orientalis* used different water sources in different seasons. According to the findings of Liu *et al*. (2016), *P. orientalis* predominantly used water from natural springs during the dry season and uptake water from shallow layers during the wet season. However, on diel time scales, *P. orientalis* used similar or identical water sources, and this could not explain the diel variation in *δ*_L,b–observed_. Our results showed that *δ*_L,b–observed_ presented a clear, dome-shaped diel pattern (Fig. 5). Midday maxima could be explained by ongoing residual leaf water enrichment from progressive loss of relatively ^18^O–depleted evaporated water. In contrast, early morning or night minima could be ascribed to the addition of ^18^O–depleted source water to leaves (Snyder *et al*., 2010). Similar diel patterns have also been observed by Welp *et al*. (2008) and Cernusak *et al*. (2005).

On the diel time scale, NSS + P performed a little better in predicting *δ*_L,b_ over ISS + P when using *α*_k–modified_ (Fig. 7a). In contrast, ISS + P performed a little better in predicting *δ*_L,b_ over NSS + P when using the previously proposed *α*_k_ value. When using same *α*_k_, differences in modeled *δ*_L,b_ assuming different isotopic conditions were not significant (*p* > 0.05); however, using different *α*_k_ values for modeled *δ*_L,b_ caused significant differences (*p* < 0.05). Using *α*_k–modified_, both ISS + P and NSS + P fit well to *δ*_L,b–observed_, with small deviations of 0.86 ± 0.57‰ and 0.66 ± 0.44‰, respectively. However, when using *α*_k–Merlivat_, ISS + P and NSS + P caused larger deviations of 2.91 ± 1.63‰ and 3.10 ± 1.43‰ from *δ*_L,b–observed_, respectively. The most significant deviations from *δ*_L,b–observed_ were caused by models using *α*_k–Cappa_ (*p <* 0.05), with deviations of 4.37 ± 2.07 (4.59 ± 1.88) ‰ for ISS+ P (NSS+ P) (Fig. 7a). It can thus be concluded that both ISS + P and NSS + P are useful for the estimation of *δ*_L,b_ at small temporal scales if *α*_k_ is well characterized.

**Figure 7.**
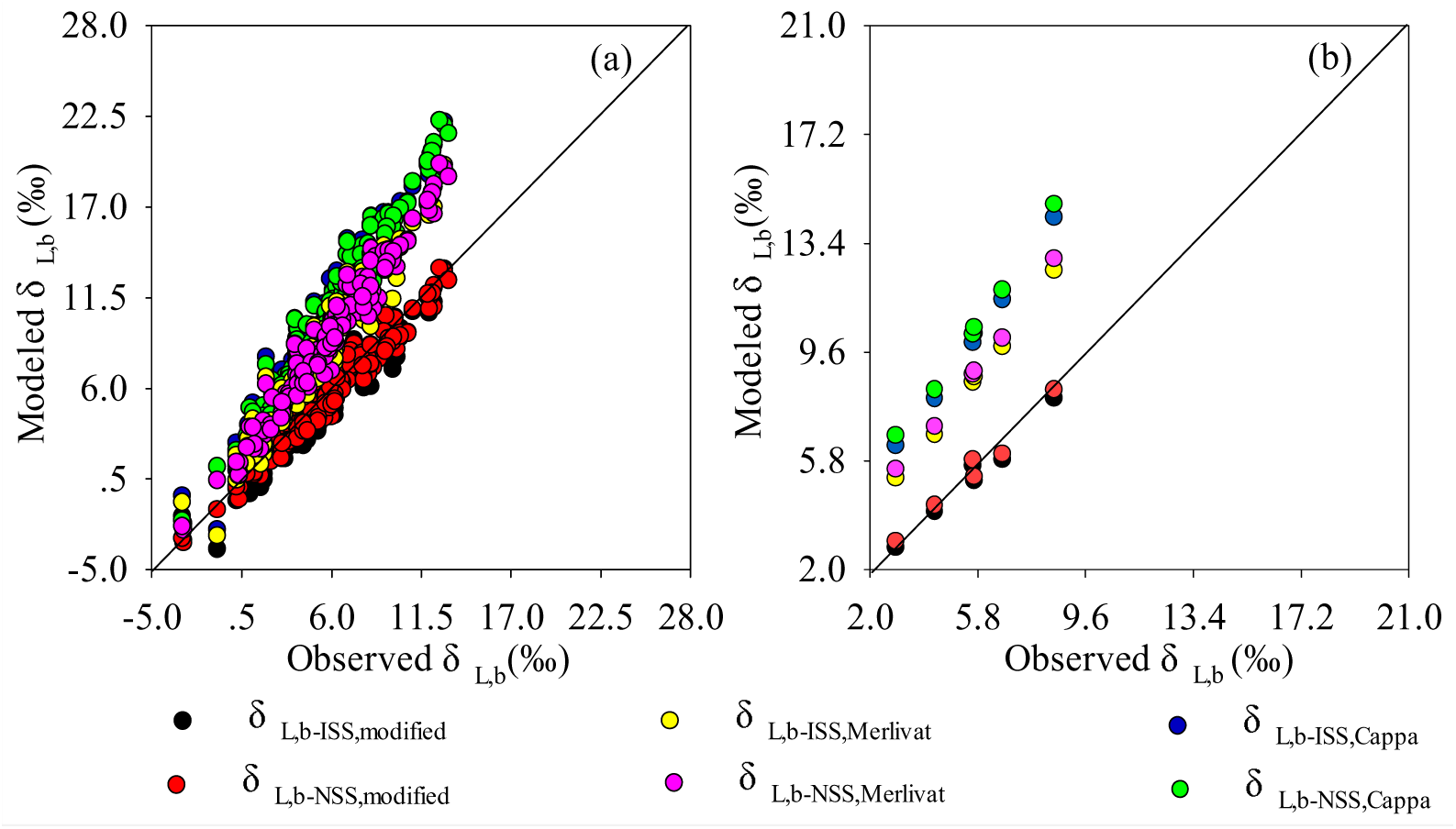
Comparison of observed and modeled *δ* O of bulk leaf water (*δ*_L,b_). Fig. 7a represents an hourly data set, while Fig. 7b is based on a daily mean data set. *δ*_L,b–ISS,Cappa_ and *δ*_L,b–ISS,Merlivat_ are simulated by ISS assumption (Eqn. 11) using *α*_k2–Cappa_ and *α*_k2–Merlivat_, respectively; *δ*_L,b–ISS,modified_ is simulated by ISS assumption (Eqn. 11) using *α*_k2–modified_; *δ*_L,b–NSS,Cappa_ and *δ*_L,b–NSS,Merlivat_ are simulated by NSS assumption (Eqn. 16) using *α*_k2–Cappa_ and *α*_k2–Merlivat_, respectively; *δ*_L,b–NSS,modified_ is simulated by NSS assumption (Eqn. 16) using *α*_k2–modified_.

On longer time scales (days), ISS + P performed better in predicting daily mean values of *δ*_L,b_ over NSS + P when using the same *α*_k_ (Fig. 7b). Using *α*_k–modified_, the ISS + P caused a negligible deviation of 0.27 ± 0.23% from daily mean values of *δ*_L,b–observed_, while NSS + P caused a larger deviation of 0.40 ± 0.30% from daily mean values of *δ*_L,b–observed_. In addition, using *α*_k–Merlivat_ and *α*_k–Cappa_, ISS + P also caused smaller deviations (2.94 ± 0.59‰ and 4.41 ± 0.85‰, respectively) from daily mean values of *δ*_L,b–observed_ than those caused by NSS + P (3.24 ± 0.63‰ and 4.73 ± 0.89‰, respectively) (Fig. 7b). This is most likely because the NSS assumption may introduce considerable complexities and uncertainties (Wang *et al*., 2014; Cernusak *et al*., 2016). Therefore, ISS + P is adequate and feasible on longer time scales.

The Péclet effect

On diel time scales, the differences between *δ*_L,b–observed_ and *δ*_L,e–ISS,modified_ or *δ*_L,e–NSS,modified_ were significant (*p* < 0.05) (Fig. 7a), while the differences between *δ*L,b–observed and *δ*L,b–ISS,modified or *δ*L,b–NSS,modified were not significant (*p* > 0.05) (Fig. 8a), indicating that the Péclet effect is required to predict plant leaf level *δ*_L,b_ at smaller temporal time scales. This is similar to the finding of Kahmen *et al*. (2008), who state that the Péclet effect is required to accurately predict leaf water enrichment in different Eucalyptus species on diurnal time scales. Ogee *et al*. (2007) also found that a spatially–explicit model was required to simulate leaf water *δ*^18^O at a high temporal resolution. It should, however, be noted that *δ*_L,e–ISS,modified_(or *δ*_L,e–NSS,modified_) rather matched *δ*L,b–observed than *δ*L,b–ISS,Cappa (or *δ*L,b–NSS,Cappa) and *δ*L,b–ISS,Merlivat (or *δ*_L,b–ISS,Merlivat_) (Figs. 7a and 8a). Clearly, considering the Péclet effect but using previously proposed *α*_k_ values may not improve model performance for the estimation of *δ*_L,b_.

**Figure 8.**
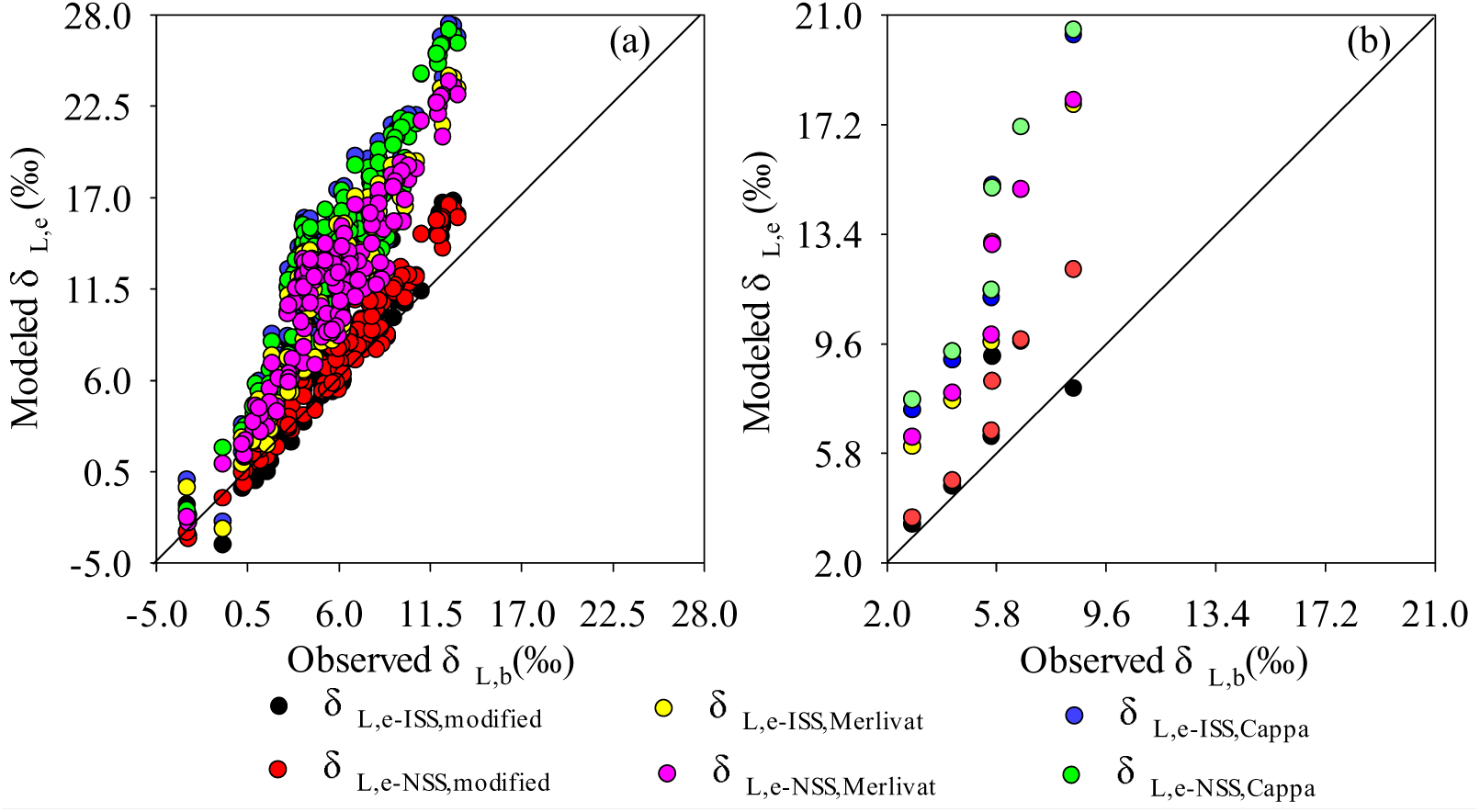
Comparison of observed *δ*^18^O of bulk leaf water (*δ*_L,b_) and modeled *δ*^18^O of leaf water at evaporative sites (*δ*_L,e_). Fig. 8(a) represents an hourly data set, while Fig. 8(b) is based on a daily mean data set. *δ*_L,e–ISS,Cappa_ and *δ*_L,e–ISS,Merlivat_ are simulated by ISS assumption (Eqn. 1) using *α*_k2–Cappa_ and *α*_k2–Merlivat_, respectively; *δ*_L,e–ISS,modified_ is simulated by ISS assumption (Eqn. 1) using *α*_k2–modified_; *δ*_L,e–NSS,Cappa_ and *δ*_L,e–NSS,Merlivat_ are simulated by NSS assumption (Eqn. 13) using *α*_k2–Cappa_ and *α*_k2–Merlivat_, respectively; *δ*_L,e–NSS,modified_ is simulated by ISS assumption (Eqn. 13) using *α*k2–modified.

On longer time scales (days), both ISS + P and NSS + P performed better in predicting daily mean values of *δ*_L,b_ over ISS and NSS assumptions without the Péclet effect when using the same *α*_k_ values (Fig. 8b), indicating that the Péclet effect is also required to predict plant leaf level *δ*_L,b_ at larger temporal scales.

Uncertainties associated with the estimation of *δ*_L,b_

Both ISS and NSS assumptions include imperfect parameterizations and uncertain parameter values. Recent methodological advances in the Off–Axis Integrated Cavity Output Spectroscopy (OA–ICOS) method have promoted the development of the water vapor isotope analyzer, which is effective for the accurate measurement of water vapor (Wang *et al*., 2009). This allows us to directly determine *δ*_V_ and *δ*_T_ (Wang *et al*., 2012; Dubbert *et al*., 2014, Song *et al*., 2015), which could greatly improve the accuracy of predictions of *δ*_L,b_. In this study, the high–frequency measurement of *δ*_V_ was expected to improve the accuracy of the estimation of *δ*_L,b_. However, the direct determination of *δ*_T_, using a laser spectrometer with an integrated gas exchange chamber (Wang *et al*., 2012), still bears uncertainties and limitations. The method developed by Wang *et al*. (2012) demands the water vapor in the chamber to be well mixed. Under such circumstance, the micro-climate conditions in the chamber are inevitably changed, that is to say, the measured flux under chamber conditions is more likely to turn into a turbulent one than under field conditions (Dubbert *et al*., 2013), thus affecting the reliability of *α*_k_ and further impacting the measured results. This effect could be more pronounced in forests with high canopies and low wind speed (Pape *et al*., 2009). In addition, the determination of *δ*_T_, using the gas-exchange chamber approach, is characterized by a poor spatial representation. Therefore, *δ*_T_ was estimated by models rather than directly measured by the gas-exchange chamber approach. Nevertheless, a good agreement was found between observed and modeled *δ*_L,b_ for both ISS + P and NSS + P when using modified *α*_k_. The parameter *α*_k_ is a key factor for determining model performance for the estimation of *δ*_L,b_. So far, there is no generally accepted calculation method for the accurate determination of *α*_k_. In our study, we tried to relate *α*_k_ with leaf water content, which is useful to estimate *δ*_L,b_. However, how *α*_k_ varies among species and with environmental changes is still not clear. Therefore, a better description of the nature of *α*_k_ under field conditions is desperately needed.

In addition, the ISS and NSS assumptions used to calculate *δ*_L,b_ on the basis of the Craig–Gordon model, in which the parameter *h*_V_ is a controlling factor at smaller temporal scales (Lee *et al*., 2007). Recent studies have suggested that high *h*_V_ could cause distortion of the computational solution (Lai *et al*., 2006; Yang *et al*., 2012). The denominator of the Craig–Gordon model is approaching zero when *h*_V_ exceeds 95%, and in such situations, outputs of models need to be excluded. In this study, *h*_V_ did not exceed 85% during the sampling period, which can improve the reliability of the Craig–Gordon model. Moreover, leaf water isotope enrichment is largely influenced by environmental and physiological parameters (Farquhar and Gan 2003;Snyder *et al*., 2010), and any variations in these parameters can be translated into *δ*_L,b_ fluctuation. Therefore, uncertainties in *δ*_L,b_ estimation might include measurement errors of environmental and physiological factors. In this study, environmental (i.e., *T*_a_, *h*_a_, *C*_v_) and physiological parameters (*g*_s_, *T*_r_, *W*, *T*_L_) were measured at a high frequency to improve the accuracy of *δ*_L,b_ prediction.

## Conclusions

In this study, *δ*_L,b_ was estimated by multiple isotopic models using various *α*_k_ values. Modeled *δ*_L,e_ was significantly larger than modeled and observed *δ*_L,b_ (*δ*_L,b–observed_) at smaller and larger temporal scales, indicating that the Péclet effect is required to predict plant leaf level *δ*_L,b_. On the diel time scale, the NSS + P performed a little better in predicting *δ*_L,b_ over the ISS + P when using *α*_k–modified_, although both ISS + P and NSS + P using *α*_k–modified_ for *δ*_L,b_ showed a good fit with *δ*_L,b–observed_. However, both ISS + P and NSS + P using previously proposed *α*_k_ (*α*_k–Cappa_ and *α*_k–Merlivat_) have significantly overestimated *δ*_L,b_ values. On longer time scale (days), ISS + P was a better estimator of daily mean values of *δ*_L,b_ than NSS + P when using the same *α*_k_ values, as the NSS assumption is likely to introduce considerable complexities and uncertainties at larger temporal scales. Also, the employment of *α*_k–modified_ led to better results in predicting daily mean values of *δ*_L,b_ than one would calculate using *α*_k–Cappa_ and *α*_k–Merlivat_. Clearly, *α*_k_ is an important controlling factor for the estimation of *δ*_L,b_. To more accurately estimate *δ*_L,b_, we call on a better description of the nature of *α*_k_.

## Acknowledgments

This study was supported by the National Natural Science Foundation of China (No.41430747), the National Science Fund for Distinguished Young Scholars (No.41401013), and the Beijing Municipal Education Commission (CEFF–PXM2018_014207_000043).

## Author Contributions Statement

Yonge Zhang designed and performed the experiment. Yonge Zhang analysed the data and wrote the manuscript. Lihua Chen, Guodong Jia contributed significantly to data analysis, manuscript preparation and practice of experiment. Xinxiao Yu revised the paper and finished the submission.

## References

Barbour MM. 2007. Stable oxygen isotope composition of plant tissue, a review. Functional Plant Biology 34, 83–94.

Cappa CD, Hendrichs MB, De Paolo DJ, Cohen RC. 2003. Isotopic fractionation of water during evaporation. Journal of Geophysical Research 108, 4525.

Cernusak LA, Farquhar GD, & Pate JS. 2005. Environmental and physiological controls over oxygen and carbon isotope composition of Tasmanian blue gum, Eucalyptus globulus. Tree Physiology 25, 129–146.

Cernusak LA, Barbour MM, Arndt SK, et al. 2016. Stable isotopes in leaf water of terrestrial plants. Plant Cell & Environment 39, 1087–1102.

Craig H, & Gordon LI. 1965. Deuterium and oxygen-18 variations in the ocean and the marine atmosphere. In Proceedings of a Conference on Stable Isotopes in Oceanographic Studies and Palaeotemperatures ed E. Tongiorgi., pp. 9–130. Lischi and Figli, Pisa.

Dongmann G, Nurnberg HW, Förstel H, Wagener K. 1974. On the enrichment of H ^18^O in the leaves of transpiring plants. Radiation and Environmental Biophysics 11, 41–52.

Dubbert M, Cuntz M, Piayda A, Werner C. 2014. Oxygen isotope signatures of transpired water vapor, the role of isotopic non–steady–state transpiration under natural conditions. New Phytologist 203, 1242–1252.

Dubbert M, Cuntz M, Piayda A, Maguás C, Werner C. 2013. Partitioning evapotranspiration–Testing the Craig and Gordon model with field measurements of oxygen isotope ratios of evaporative fluxes. Journal of Hydrology 49614, 142–153.

Farquhar GD, & Gan KS. 2003. On the progressive enrichment of the oxygen isotopic composition of water along leaves. Plant, Cell and Environment 26, 801–819.

Farquhar GD, & Cernusak LA. 2005. On the isotopic composition of leaf water in the non–steady state. Functional Plant Biology 32, 293–303.

Farquhar GD, Cernusak LA, Barnes B. 2007. Heavy water fractionation during transpiration. Plant Physiology 143, 11–18.

Ferrio JP, Pou A, Florez–Sarasa I, Gessler A, Kodama N, Flexas J. & Ribas–Carbo, M. 2012. The Peclet effect on leaf water enrichment correlates with leaf hydraulic conductance and mesophyll conductance for CO_2_. Plant Cell and Environment 35, 611–625.

Gessler A, Ferrio JP, Hommel R, Treydte K, Werner RA, & Monson RK. 2014. Stable isotopes in tree rings, towards a mechanistic understanding of isotope fractionation and mixing processes from the leaves to the wood. Tree Physiology 34, 796–818.

Gonantini R, Gratziu S, & Tongiorgi E. 1965. Oxygen isotope composition of water in leaves. In Isotopes and Radiation in Soil-Plant Nutrition Studies, pp. 405–410. IAEA, Vienna.

Hoffmann G, Cuntz M, Weber C, et al. 2004. A model of the Earth’s Dole effect. Global Biogeochemical Cycles 18, 1008.

Kahmen K, Simonin KP, Tu KP, Merchant A, Callister A, Siegwolf R, Dawson TE, Arndt SK. 2008. Effects of environmental parameters, leaf physiological properties and leaf water relations on leaf water delta ^18^O enrichment in different Eucalyptus species. Plant Cell and Environment 31, 738–751.

Lai CT, Ehleringer JR, Bond BJ, Paw KTU. 2006. Contributions of evaporation, isotopic non–steady state transpiration and atmospheric mixing on the δ18O of water vapour in Pacific Northwest coniferous forests. Plant Cell and Environment 29, 77–94.

Lee XH, Kim K, Smith R. 2007. Temporal variations of the ^18^O/^16^O signal of the whole-canopy transpiration in a temperate forest. Global Biogeochemical Cycles 21, 3013.

Lee X, Griffis TJ, Baker JM, Billmark KA, Kim K, Welp LR. 2009. Canopy-scale kinetic fractionation of atmospheric carbon dioxide and water vapor isotopes. Global Biogeochemical Cycles 23, 1002.

Liu ZQ, Yu XX, Jia GD, Jia JB, Lou YH, Lu WW. 2017. Contrasting water sources of evergreen and deciduous tree species in rocky mountain area of Beijing, China. Catena 150, 108–115.

Luo L, Yu WS, Wan SM, Zhou P. 2013. Advances in the study of stable isotope composition of leaf water in plants. Acta ecologica sinica 33, 1031–1041.

Luz B, Barkan E. 2011. The isotopic composition of atmospheric oxygen. Global Biogeochemical Cycles 25, 3001.

Mathieu R, Bariac T. 1996. A numerical model for the simulation of stable isotope profiles in drying soils. Journal of Geophysical Research Atmospheres 101, 12685–12696.

Merlivat L. 1978. Molecular diffusivities of H_2_^16^O, HD^16^O, and H ^18^O in gases. Journal of Chemical Physics 69, 2864–2871.

Ogée J, Cuntz M, Peylin P, Bariac T. 2007. Non-steadystate, non-uniform transpiration rate and leaf anatomy effects on the progressive stable isotope enrichment of leaf water along monocot leaves. Plant, Cell and Environment 30, 367–387.

Pape L, Ammann C, Nyfeler–Brunner A, Spirig C, Hens K, Meixner FX 2009. An automated dynamic chamber system for surface exchange measurement of non–reactive and reactive trace gases of grassland ecosystems. Biogeosciences 6, 405–429.

Rothfuss Y, Braud I, Moine LN, Biron P, Durand JL, Vauclin M, Bariac T. 2012. Factors controlling the isotopic partitioning between soil evaporation and plant transpiration, assessment using a multi–objective calibration of Si SPAT–Isotope under controlled conditions. Journal of hydrology 442, 75–88.

Snyder KA, Monnar R, Poulson SR, Hartsough P, Biondi F. 2010. Diurnal variations of needle water isotopic ratios in two pine species. Trees 24, 585–595.

Song X, Barbour MM, Farquhar GD, Vann DR, Helliker BR. 2013. Transpiration rate relates to within–and across–species variations in effective path length in a leaf water model of oxygen isotope enrichment. Plant Cell and Environment 36, 1338–1351.

Song X, Loucos KE, Simonin KA, Farquhar GD, Barbour MM. 2015. Measurements of transpiration isotopologues and leaf water to assess enrichment models in cotton. New Phytologist 206, 637–646.

Stewart MK. 1975. Stable isotope fractionation due to evaporation and isotopic exchange of falling water drops. Application to atmospheric processes and evaporation of lakes. Journal of Geophysical Research 80, 1133–1146.

Sullivan PF, Welker JM. 2007. Variation in leaf physiology of Salix arctica within and across ecosystems in the High Arctic,test of a dual isotope Delta ^13^C and Delta ^18^O. conceptual model. Oecologia 151, 372–386.

Sun SJ, Meng P, Zhang JS, Wan X, Zheng N, He CX. 2014. Partitioning oak woodland evapotranspiration in the rocky mountainous area of North China was disturbed by foreign vapor, as estimated based on non–steady. Agricultural and Forest Meteorology 184, 36–47.

Walker, CD, Brunel, JP, Dighton J, Holland K, Leaney F, Mc EwanK, Mensforth L, Thorburn P, Walker C. 2001. The use of stable isotopes of water for determining sources of water for plant transpiration. In, Unkovich M, Pate J, McNeil A, Gibbs JD eds. Stable isotope techniques in the study of biological processes and functioning of ecosystems. Kluwer Academic Press, Dordrecht, pp 57–89.

Wang L, Good SP, Caylor KK, Cernusak LA. 2012. Direct quantification of leaf transpiration isotopic composition. Agricultural and Forest Meteorology 154–155, 127–135.

Wang P, Yamanaka T, Li XY, Wei Z. 2015. Partitioning evapotranspiration in a temperate grassland ecosystem, Numerical modeling with isotopic tracers. Agricultural and Forest Meteorology 208, 16–31.

Wang L, Caylor, KK, Dragoni, D. 2009. On the calibration of continuous, high-precision δ18O and δ2H measurements using an off–axis integrated cavity output spectrometer. Rapid Commun. Mass Spectrom 23, 530–536.

Welp LR, L X, Kim K, Griffis TJ, Billmark KA, Baker JM. 2010. δ18O of water vapour, evapotranspiration and the sites of leaf water evaporation in a soybean canopy[J]. Plant Cell & Environment 319, 1214–1228.

Welp LR, Keeling RF, Meijer HAJ, et al. 2011. Interannual variability in the oxygen isotopes of atmospheric CO2 driven by El Nino. Nature 477, 579–582.

Wen XF, Zhang SC, Sun XM, Yu GR. 2008. Recent advances in H_2_O enrichment in leaf water. Journal of Plant Ecology 32, 961–966.

Xiao Q, Ye WJ, Zhu Z, Chen Y, Zheng HL. 2005. A simple non–destructive method to measure leaf area using digital camera and Photoshop software. Chinese Journal of Ecology 24, 711–714.

Yang B, Xie FT, Wen XF, Sun XM, Wang JL. 2012. Diurnal variations of soil evaporation δ18O and factors affecting it in cropland in North China. Chinese Journal of Plant Ecology 366, 539–549.

Yepez EA, Williams DG, Scott RL, Lin G. 2003. Partitioning overstory and understory evapotranspiration in a semiarid savanna woodland from the isotopic composition of water vapor. Agricultural and Forest Meteorology 119, 53–68.

Yepez EA, Huxman TE, Ignace DD, English NB, Weltzin JF, Castellanose AE, Williams DG. 2005. Dynamics of transpiration and evaporation following a moisture pulse in semiarid grassland, A chamber–based isotope method for partitioning flux components. Agricultural and Forest Meteorology 132, 359–376.

Yu GR, Wang QF. 2010. Ecophysiology of plant photosynthesis, Transpiration, and Water Use, ed. Li Y, Chen SS, Science press, Beijing.

Zhang Y, Yu X, Chen L, Jia GD. 2018. Comparison of the partitioning of evapotranspiration – numerical modeling with different isotopic models using various kinetic fractionation coefficients[J]. Plant & Soil 14, 1–22.

Zhou Y, Grice K, Chikaraishi Y, Stuartwilliams H, Farquhar GD, Ohkouchi N. 2011. Temperature effect on leaf water deuterium enrichment and isotopic fractionation during leaf lipid biosynthesis, results from controlled growth of C_3_ and C_4_ land plants. Phytochemistry 72, 207–213.

